# Nanoprojectile Secondary Ion Mass Spectrometry Enables Multiplexed Analysis of Individual Hepatic Extracellular Vesicles

**DOI:** 10.1101/2023.08.21.554053

**Authors:** Seonhwa Lee, Dmitriy S. Verkhoturov, Michael J. Eller, Stanislav V. Verkhoturov, Michael A. Shaw, Kihak Gwon, Yohan Kim, Fabrice Lucien, Harmeet Malhi, Alexander Revzin, Emile A. Schweikert

## Abstract

Extracellular vesicles (EVs) are nanoscale lipid bilayer particles secreted by cells. EVs may carry markers of the tissue of origin and its disease state which makes them incredibly promising for disease diagnosis and surveillance. While the armamentarium of EV analysis technologies is rapidly expanding, there remains a strong need for multiparametric analysis with single EV resolution. Nanoprojectile (NP) secondary ion mass spectrometry (NP-SIMS) relies on bombarding a substrate of interest with individual gold NPs resolved in time and space. Each projectile creates an impact crater of 10−20 nm in diameter while molecules emitted from each impact are mass analyzed and recorded as individual mass spectra. We demonstrate the utility of NP-SIMS for analysis of single EVs derived from normal liver cells (hepatocytes) and liver cancer cells. EVs were captured on antibody (Ab)-functionalized gold substrate then labeled with Abs carrying lanthanide (Ln) MS tags (Ab@Ln). These tags targeted four markers selected for identifying all EVs, and specific to hepatocytes or liver cancer. NP-SIMS was used to detect Ab@Ln-tags co-localized on the same EV and to construct scatter plots of surface marker expression for thousands of EVs with the capability of categorizing individual EVs. Additionally, NP-SIMS revealed information about the chemical nano-environment where targeted moieties co-localized. Our approach allowed analysis of population heterogeneity with single EV resolution and distinguishing between hepatocyte and liver cancer EVs based on surface marker expression. NP-SIMS holds considerable promise for multiplexed analysis of single EVs and may become a valuable tool for identifying and validating EV biomarkers of cancer and other diseases.

## INTRODUCTION

Cancer remains a major cause of death in the US and the world. Several cancer types including hepatocellular carcinoma (HCC) are detected at an advanced or metastatic stage and have low 5-year survival rates.^1–3^ There is a need to identify biomarkers for early cancer detection and for monitoring response to therapy.^4–6^ Tumor-derived markers present in blood or other physiological fluids may be used for cancer surveillance. The analysis of cancer biomarkers in a physiological fluid, termed liquid biopsy, may focus on circulating tumor cells (CTCs),^7, 8^ cell-free DNA (cf)DNA,^9, 10^ proteins,^11^ and extracellular vesicles (EVs)^12–14^. EVs hold particular promise for cancer surveillance because these nanoparticles (50−1000 nm in diameter) may be shed by the tumor into circulation and help identify the tumor type and its pathological state.^12–14^ However, EVs released by the tissue of interest represent a small fraction of total EVs (<1%) in a biological fluid such as blood. This motivates the need for sensitive and specific technologies that analyze EV heterogeneity to profile correlates of disease progression or therapy response.

Technologies for detecting EVs may be broadly placed into three categories: 1) immunoassays, 2) mass spectrometry (MS), and 3) single particle analysis.

While immunoassays (e.g. ELISAs) may be used to quantify expression of EV biomarkers, they do not offer the ability to associate multiple markers with the same EV or the same subset of EVs.

Mass spectrometry approaches including, liquid chromatography (LC)-MS/MS, matrix assisted laser desorption (MALDI)-MS, and inductive coupled plasma (ICP)-MS characterize EVs en masse, report the average chemical composition of a population and do not offer single EV resolution.^15–18^ While there is MS technology specifically developed for the analysis of single cells (cytometry by time of flight – CyToF),^19^ it does not have sufficient sensitivity for detecting individual EVs, and no reports exist of CyToF used for analysis of single EVs.

Single EV analysis may be accomplished using several approaches including flow cytometry, Raman spectroscopy and optical trapping.^20–23^ These methods have allowed analysis of the size, shape, content, and biological activity of individual EVs. Flow cytometry is arguably the most commonly used approach for quantitative and compositional analysis of single EVs.^24, 25^ It has been used, for example, to characterize EV subsets produced by colorectal cancer cell lines and platelets.^25–29^ However, flow cytometry and other single EV analysis approaches provide only limited information on the relative abundancies of biomarkers. Despite the recent progress, there remains a need for technologies for high-throughput and multiplexed analysis of single EVs that also allow quantification of EV marker expression.

We reasoned that a surface analysis technology, nanoprojectile (NP) secondary ion mass spectrometry (NP-SIMS), could be an effective detection modality that enables sensitive and multiplexed characterization of individual EVs. This technology is unique amongst the tools currently available for EV analysis in that it performs mass spectrometry analysis on individual EVs and combines single EV resolution with quantitative analysis of surface marker expression. Our team has pioneered the SIMS approach whereby a surface is stochastically bombarded with a suite of individual nanoprojectiles (e.g. ∼1 keV/atom Au ^4+^) in the event-by-event bombardment detection mode.^30–33^ A bombardment event generates a crater of ∼10 nm in diameter within an organic layer on the surface; ionized ejecta from each bombardment event are detected and mass spectra from millions of such events are compiled. This approach has been used by us previously to examine the heterogeneity of biointerfaces,^34, 35^ thin films,^36, 37^ nanoparticles,^38–40^ and integrated circuits. We have recently demonstrated the feasibility of NP-SIMS for EV analysis using antibodies (Abs)-labeled with halide MS tags and targeting abundant and ubiquitous surface markers, CD63 and CD81.^41^

We wanted to further establish NP-SIMS as a method for EV analysis by identifying tagging strategies better suited for multiplexing and by focusing on disease-specific EV surface markers. We describe the use of NP-SIMS in combination with lanthanide (Ln) tags for multiplexed analysis of individual EVs. To prove this concept, EVs were isolated from hepatic cells and labeled with Ln-tagged Abs against four surface markers that may be grouped into categories of conserved EV marker (CD63), hepatic (CYP2E1) and cancer (EpCAM, AFP). NP-SIMS analysis allowed detection of these surface markers from individual EVs and could be used to differentiate cancer and normal hepatic EVs based on the type and levels of surface marker expression.

## RESULTS AND DISCUSSION

### NP-SIMS as a strategy for multiplexed analysis of individual EVs

NP-SIMS is used to analyze the chemical composition of nanoobjects immobilized on a conductive surface. For our experiments, EVs were captured on Au substrates functionalized with Abs targeting generic surface marker CD63. The captured EVs were labeled with Ab@Ln tags targeting generic (CD63), hepatic (CYP2E1) and cancer (EpCAM, AFP) surface markers (see **Figure 1**). These EV samples were then analyzed using NP-SIMS to detect the presence and quantify levels of surface markers. NP-SIMS is distinct from conventional SIMS in the character of the projectile, a gold nanoparticle (Au_400_^4+^) of ∼1 keV/atom kinetic energy, and in the mode of bombardment which is at the level of one Au ^4+^ at any time. The individual impacts are repeated at a rate of ∼1 kHz.

**Figure 1.**
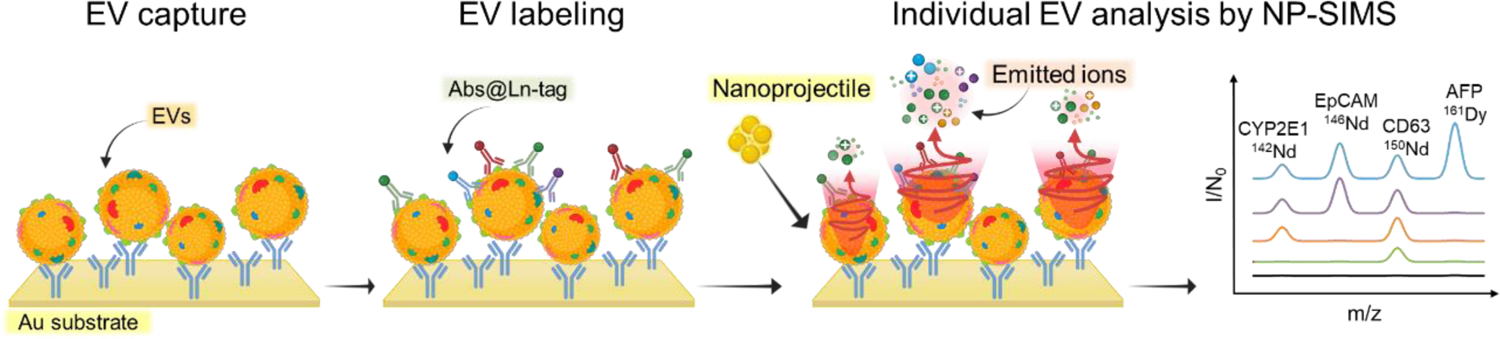
Probing composition of single EVs using NP-SIMS. EVs were captured on the Au surface, labeled with Abs carrying Ln-tags and then analyzed by NP-SIMS. Individual EVs were analyzed for surface marker expression.

Interaction of one projectile with the substrate creates a 10 nm crater^31^ and produces ejecta in an amount sufficient to be detected by the instrument. We typically use 10^6–7^ projectiles to probe the surface, each projectile separated in time as noted already, and each projectile impact producing a mass spectrum. Therefore, 10^6–7^ individual mass spectra containing information on co-localized moieties are collected from each analysis run.

How do we ensure that individual EVs are analyzed? First, we optimized EV capture conditions to produce surfaces with ∼5% EV surface coverage as determined by SEM and fluorescence microscopy (**Figure S1**). This decreases the possibility of a single impact affecting two or more EVs. Second, we use a relatively small number of projectiles (10^6–7^) that strike the substrate randomly and analyze ∼0.3% of the surface. This makes it statistically unlikely that two or more impacts analyze the same region of the surface (probability < 3 × 10^-6^).

### Assessing EV labeling with Ab@Ln-Tags

Prior to MS analysis, we wanted to validate strategies for capture and labeling of EVs. We used SPR for this purpose because of the similarity in surface composition between SPR and NP-SIMS chips and the ability to monitor EV-tag interactions in real-time without additional labeling.^42^ Au SPR chips were functionalized with carboxyl-terminated alkanethiols and then activated with EDC/NHS for attachment of anti-CD63 Abs. The surfaces were also blocked with BSA to minimize non-specific binding (**see Figure S2**). For EV labeling, we used tags with unique Ab-MS label pairings: anti-CD63@^150^Nd, anti-CYP2E1@^142^Nd, anti-EpCAM@^146^Nd, and anti-AFP@^161^Dy (see Materials and Method for details).^43^ These tags were introduced individually and sequentially into the SPR instrument to characterize interactions with captured EVs as illustrated in **Figure 2A**. A typical SPR sensorgram may be seen in **Figure 2B**. It shows successful capture of EVs followed by binding of Ab@Ln tag molecules. The change in SPR response (RU) resulting from binding of Ab@Ln to the EVs was quantified and used to compare target marker expression levels. These results, presented in **Figure 2B,C**, demonstrated differences in the expression of the four surface markers. It is of interest that cancer markers, EpCAM and AFP,^12, 44, 45^ were observed to a have higher expression level compared to normal hepatic marker CYP2E1. This observation is logical given that EVs used for analysis were harvested from media conditioned by the liver cancer cell line (HepG2 cells). We carried out a series of experiments to assess non-specific interactions using SPR (see **Figure S3**). When captured EVs were labeled with anti-CD63@^150^Nd, SPR signal was significantly higher (32 RU) than that produced by the anti-IgG (isotype control)@^150^Nd (< 3 RU). When a solution of anti-CD63@^150^Nd tag interacted with a SPR chip that did not have EVs, the signal was similar to that of the isotype control (< 3 RU). These experiments confirmed that tags interacted specifically with EVs and that the presence of target-specific Abs was necessary for EV/tag interaction to take place. These findings support the specificity of interactions between Abs@Ln-tag and EVs, which is important for accurate and reliable detection and analysis of EVs in biological samples.

**Figure 2.**
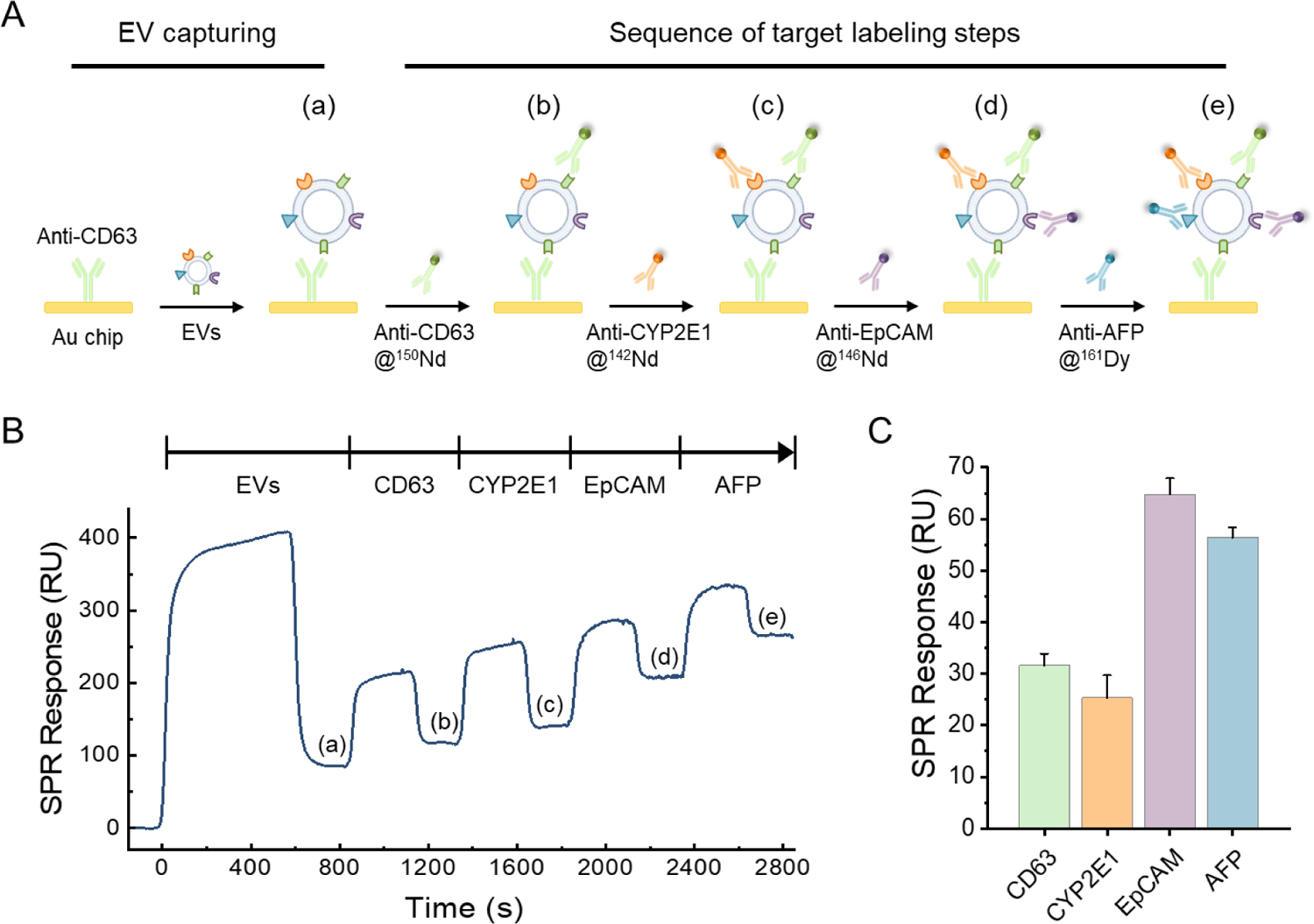
Using SPR to characterize EV capture and labeling. (A) Cartoon describing experiment setup. HepG2 EVs are first captured on an SPR chip and then sequentially labeled with Ab@Ln tags. (B) A typical SPR sensogram where signal (a) is due to binding of EVs while signals (b) thru (e) arise from binding of Ab@Ln tags. (C) SPR signals for binding of four different Ab@Ln tags investigated in this study. The SPR response in resonance units (RU) was determined from the baseline change before and after injecting each Ab@Ln tag. Data are displayed as mean ± SD (n=3).

Does the sequence of the EV labeling steps matter? Given our interest in multiplexed detection of EV surface markers, we wanted to optimize the labeling process focusing on the potential steric hindrance when using different Ab-tag conjugates. We proceeded to vary the sequence of labeling EVs using the four tags of interest while assessing SPR responses. As may be appreciated from **Figure S4**, labeling for more abundantly expressed EpCAM first resulted in much weaker signals for AFP, CD63 and CYP2E1 labeled second, third and fourth, respectively (see **Figure S4B**). Conversely, starting the sequence of labeling steps with CD63, followed by CYP2E1, EpCAM and AFP produced stronger signals for underabundant surface markers like CYP2E1 (see **Figure 2C**). These observations informed the sequence of labeling steps when preparing samples for NP-SIMS analysis and underscored the need to carefully examine labeling protocols.

### Detecting individual EVs using NP-SIMS

Having optimized EV labeling and confirmed specificity of EV-tag interactions with SPR, we proceeded to analyze the chemical composition of tagged EVs with NP-SIMS. Four different Ln ions ^150^Nd, ^142^Nd, ^146^Nd, and ^161^Dy were paired with Abs to create MS tags targeting EV surface markers of interest.

At the first stage of characterization, we prepared EV-containing substrates labeled with either one, two, three or four Ab@Ln-tags; 1-tag (anti-CD63@^150^Nd tag), 2-tag (anti-CD63@^150^Nd and anti-CYP2E1@^142^Nd), 3-tag (anti-CD63@^150^Nd, anti-CYP2E1@^142^Nd, and anti-EpCAM@^146^Nd), and 4-tag (anti-CD63@^150^Nd, anti-CYP2E1@^142^Nd, anti-EpCAM@^146^Nd, and anti-AFP@^161^Dy). A sample of EVs without tags was used as the negative control. It is worth mentioning that functionalization of Au substrates for NP-SIMS analysis follows protocols similar to those described in SPR characterization; Au surfaces were treated with carboxyl terminated thiols, activated by EDC/NHS, and incubated with anti-CD63 Abs.

The mass spectra of characteristic ions associated with four Ln-tags were identified and compared in **Figure S5**. We first examined EVs tagged with anti-CD63@^150^Nd and observed the characteristic peaks for ^150^Nd^+^ and ^150^NdO^+^. The measured yield for tag related ions was found to be 1.3 percent. Note that the measured abundance of a tag will depend on the fraction of the surface containing EVs. Similarly, other tagged EV surfaces (i.e., 2-tag, and 3-tag) showed abundant emission of ions for a given Ln-tag carrying Abs against the target marker. Conversely, mass spectrometry signals for Ln ions were not observed in the non-tagged EV surface. Importantly, after the labeling of EVs with the four tags, we detected characteristic ions associated with all four tags, including ^150^Nd^+^ and ^150^NdO^+^ for CD63, ^142^Nd^+^ and ^142^NdO^+^ for CYP2E1, ^146^Nd^+^ and ^146^NdO^+^ for EpCAM and ^161^Dy^+^ and ^161^DyO^+^ for AFP. These results clearly demonstrate the analytical capacity of NP-SIMS for detecting up to four co-localized Abs from the individual EVs with enhanced signals.

To further evaluate the specificity of NP-SIMS, we compared the Ln-tag signal from EVs exposed to isotype control Abs vs. target-specific Abs (see **Figure S6**). EVs produced by HepG2 were labeled with either anti-CD63@^150^Nd-tag or isotype control, anti-rabbit IgG@^150^Nd, and analyzed by NP-SIMS. As shown in **Figure S6**, Nd mass peaks were only observed for tags targeting CD63. Another negative control was a substrate functionalized with anti-CD63 but without EVs. Incubating this substrate with anti-CD63@^150^Nd produced signals similar to isotype control tags incubated (see **Figure S6)** suggesting minimal nonspecific interactions between Ab@Ln tag and the Au substrate without EVs. These findings once again highlight the specificity of the NP-SIMS assay towards EV surface markers.

### Comparing EVs from normal cells and cancer cells

After validating the Ab@Ln tagging strategy, we carried out NP-SIMS analysis of EVs from healthy primary human hepatocytes and a hepatoblastoma cell line (HepG2 cells). Both cell types were cultured in EV-free DMEM to eliminate the confounding effect of EVs endogenous to serum. Nanoparticle tracking analysis (NTA) revealed that EVs secreted by both cell types had similar size distribution, ranging from 50 to 300 nm with an average diameter of 120 nm (see **Figure S7**). The isolated EVs were resuspended in PBS at 10^9^ particles/mL and were used for all NP-SIMS analysis experiments. Our ultimate goal is to leverage NP-SIMS for identifying EV-based biomarkers and signatures of HCC and other liver diseases. As a step toward this goal, we explored expression of four surface markers relevant to liver diseases and cancer; CD63 (conserved EV marker), CYP2E1 (hepatic marker), EpCAM (epithelial marker/cancer marker), and AFP (liver cancer marker).^12, 44–46^

EVs were captured on Au substrates and labeled with Ab@Ln tags as described above. Untagged EVs were used as the control. The mass spectra of tagged and non-tagged EVs are presented in **Figure 3A**. The abundant emission of ^150^Nd^+^ and ^150^NdO^+^ for CD63, ^142^Nd^+^ and ^142^NdO^+^ for CYP2E1, ^146^Nd^+^ and ^146^NdO^+^ for EpCAM and ^161^Dy^+^ and ^161^DyO^+^ for AFP was observed from both normal and cancer EVs, but not from control EVs without tags. We next compared the level of surface marker expression in cancer and normal EVs. To quantify EV markers, we developed a methodology that uses ions characteristic of the target object or the substrate.^34, 39^ We monitored four co-localized Ab@Ln-tags and determined a relative abundance (%) of each tag which correlates to surface expression of the target protein. As seen from **Figure 3B**, the analysis of relative abundance (%) revealed that normal EVs had a higher level of CYP2E1 (hepatic marker) concomitant with lower levels of AFP (liver cancer marker) and EpCAM (generic cancer). In cancer EVs, levels of AFP increased whereas levels of CYP2E1 marker decreased. These observations are consistent with the notion that, compared to normal liver tissue, liver cancer tissue is less differentiated with lower expression of mature hepatic markers (e.g. CYP2E1) and upregulated expression of fetal markers like AFP.^47^

**Figure 3.**
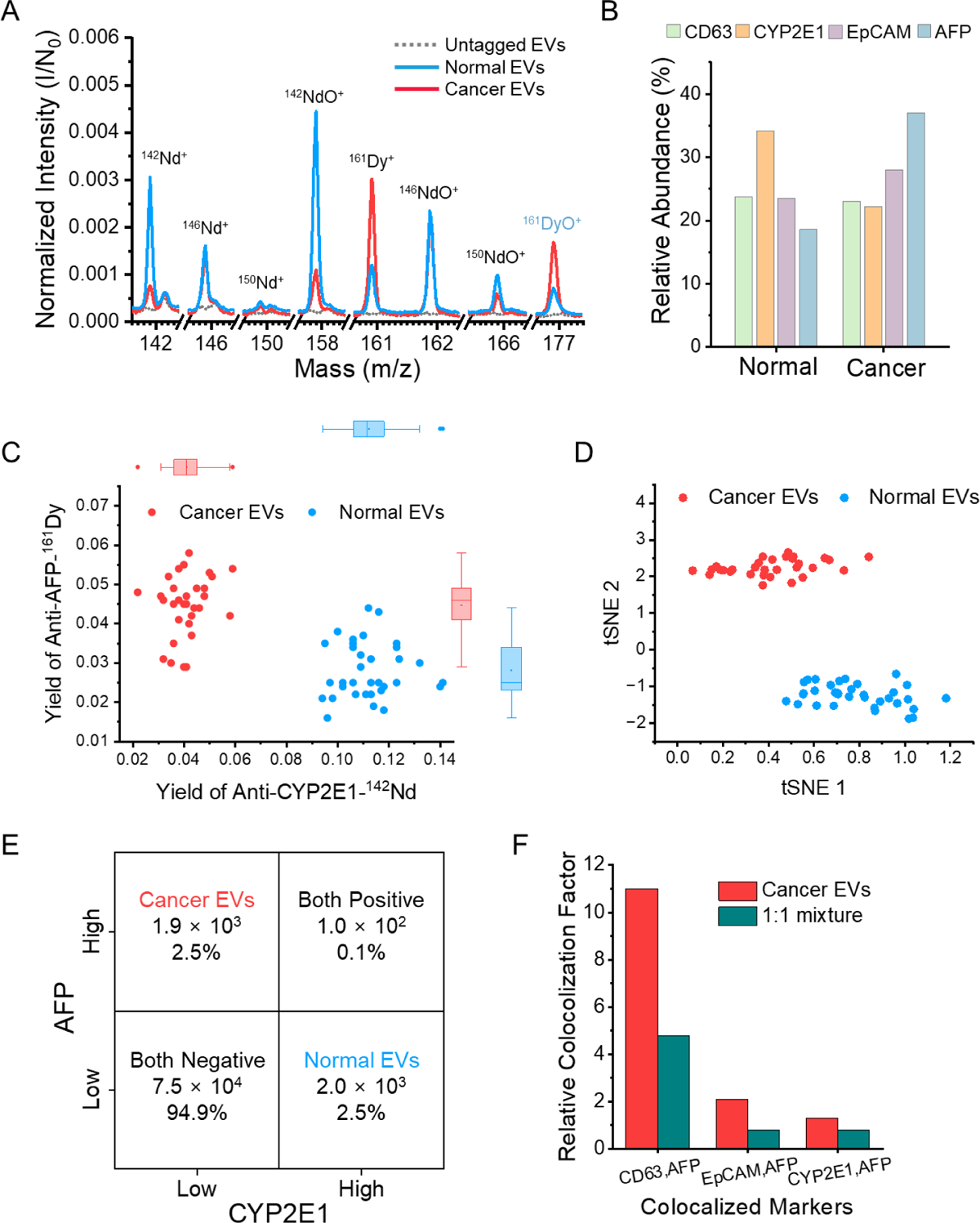
NP-SIMS analysis of EVs from normal and malignant liver cells. (A) Selected region of mass spectra comparing tagged and untagged EVs. Note that signals originate from both normal and cancer EVs labeled with Abs@Ln tags but not from EVs without tags. The data are binned to the nearest 0.02 m/z for comparison. (B) Comparison of relative abundance (%) of the tags (surface marker expression) on EVs produced by normal hepatocytes and by cancer cells. Note that cancer marker AFP is expressed higher in cancer EVs while marker of well-differentiated hepatocytes CYP2E1 is higher in normal EVs. (C,D) Analysis of EV heterogeneity in biomarker expression. Scatter plot representation of EV analysis based on expression of AFP and CYP2E. tSNE scores based on abundance of AFP, CYP2E1, and EpCAM. (E) Analysis of 1:1 mixture where individual EVs are identified as originating from HepG2 or normal cells. (F) Relative colocalization factors, *CF*, for APF colocalized with the other three markers, CD63, EpCAM, CYP2E1.

Overall, **Figure 3A,B** confirmed that (1) detected ions are specific to the target of interest and have variable levels of expression and (2) levels of CYP2E1 and AFP trend in the opposite directions for normal and liver cancer EVs.

### Characterizing EV heterogeneity

NP-SIMS provides information on both molecular abundance and EV heterogeneity. To highlight this point, we arranged output from 10^6^ mass spectra in a scatter plot format as shown in **Figure 3C**. Given the large number of mass spectra collected, each dot represents 1000 EVs. A determination of whether a nanoprojectile landed on an EV or the underlying substrate was made based on the presence of CD63 signal (^150^Nd) in the mass spectrum. Thus, events in **Figure 3C** were “gated” for CD63 expression and report on co-localization of CD63 with AFP and CYP2E1. As seen from these data (see **Figure 3C and S8**), NP-SIMS allows us to analyze EV heterogeneity, and to distinguish normal vs. cancer EVs based on the levels of surface markers, CYP2E1 and AFP. The analysis of NP-SIMS data can be extended to encompass the abundance and co-localization of multiple biomarkers using multivariate analysis tools such as t-distributed stochastic neighbor embedding, t-SNE. This is a powerful data analysis tool used in mass spectrometry to aid in biomarker discovery and data interpretation.^48, 49^ **Figure 3D** highlights that t-SNE analysis of multiple biomarkers detected by NP-SIMS (see **Figure 3A**) differentiates EVs harvested from normal hepatocytes and HepG2 cells.

### Establishing a data analysis method for characterizing individual EVs with NP-SIMS

The section above describes a strategy for binning and averaging NP-SIMS measurements from 1000 EVs to distinguish between normal and HepG2 samples. As shown in **Figure 3D**, such data are amenable to multivariate analysis and can form the basis of a data library for identifying the source of EVs. However, averaging, even at the level of 1000 EVs, may be suboptimal for analysis of clinical samples containing a mixture of EV types. In fact, averaging approaches could not be used to correctly predict the composition of the sample containing a 1:1 mixture of cancer and normal EVs. Therefore, we wanted to develop a data analysis strategy to count signals from single EVs and avoid ensemble averaging entirely. NP-SIMS data collection occurs in a pulse counting mode, where secondary ions are detected one-by-one with a microchannel plate-based multi-anode detector (described further elsewhere).^31^ Using this detection scheme, up to eight ions of the same mass to charge can be detected from a single projectile impact, thus up to eight tag specific ions (e.g. ^150^Nd^+^) can be detected from a single EV. As previously discussed, AFP expression is upregulated in EVs from HepG2 cells, while CYP2E1 is downregulated. In contrast, EVs from normal cells display low levels of AFP and higher levels of CYP2E1. We gated data based on the presence of anti-CD63 (detection of ^150^Nd^+^ or ^150^NdO^+^) to ensure that EVs were being analyzed and then grouped measurements from individual EVs based on the number of detected AFP and CYP2E1 tag ions**. Figure S9** compares the probability of detecting different combinations of AFP and CYP2E1 tag ions ranging from zero to three ions of each tag. HepG2 EVs were characterized by the detection of one to three AFP tag ions and zero CYP2E1 tag ions. While normal EVs were more likely to result in the detection of one to three CYP2E1 tag ions with zero AFP tag ions or two CYP2E1 tag ions with one AFP tag. Both samples displayed similar probabilities for the detection of zero or one ion related to each tag. These results are consistent with our previous findings where AFP and CYP2E1 displayed an inverse relationship for EVs collected from HepG2 cells and normal hepatocytes.

When applied to the analysis of a 1:1 mixture of HepG2 and normal EVs, this methodology of ion counting allowed us to accurately detect the composition of EVs captured on the substrate. As shown in **Figure 3E**, 2.0 × 10^3^ and 1.9 × 10^3^ EVs were grouped into normal and cancer categories, respectively. Thus, we demonstrated that NP-SIMS in combination with the data analysis strategy may be used to correctly predict the composition of a sample containing a mixture of EVs.

### Examining the chemical nano-environment of normal and cancer EVs

As described previously, NP-SIMS allows for detection of colocalized species. We posited that colocalization of surface markers may be non-random and different for normal versus HepG2 EVs. To enable this analysis, we established colocalization factors (CF) for individual pairs of Ln tags. Methodology for computing CF values may be found in Supporting Information (**Figure S10**). Briefly, CF value was determined by dividing the number of co-detection events for two characteristic ions/markers (e.g. CD63 and AFP) by the number of detection events for all four markers. Determination of colocalization of two Ab@Ln tags is based on the size of the probing area (diameter ∼10 nm) and the size of the tagged area of a biomarker (∼15 nm). If the distance between the tagged areas exceeds the diameter of the probing area, the markers are considered non-colocalized. Comparison of CF values for all four surface markers revealed that CD63 and AFP appeared together non-randomly, were co-localized in the case of HepG2 EVs but not normal EVs (see **Figure 3F**). Relative CFs, defined as a ratio of cancer to normal EVs, were ∼11 for HepG2 EVs and 4.8 for the 1:1 mixture population of HepG2 and normal EVs. Therefore, the co-localization analysis approach offered another way to determine composition of a mixed population of EVs. In addition, these results highlight the fact that NP-SIMS data not only inform on the relative abundance of molecules but also reveal how these molecules co-localize on a nanometer scale. It may be possible and beneficial to use both parameters as biomarkers of disease in the future.

### Benchmarking NP-SIMS analysis against SPR and nanoflow cytometry

We wanted to rigorously compare NP-SIMS results against established methods, SPR and nanoflow cytometry. As described in **Figure 4A**, EVs were harvested from media conditioned by liver cancer hepatoblastoma-derived cells (HepG2) and were analyzed by NP-SIMS and either SPR or nanoflow cytometry.

**Figure 4.**
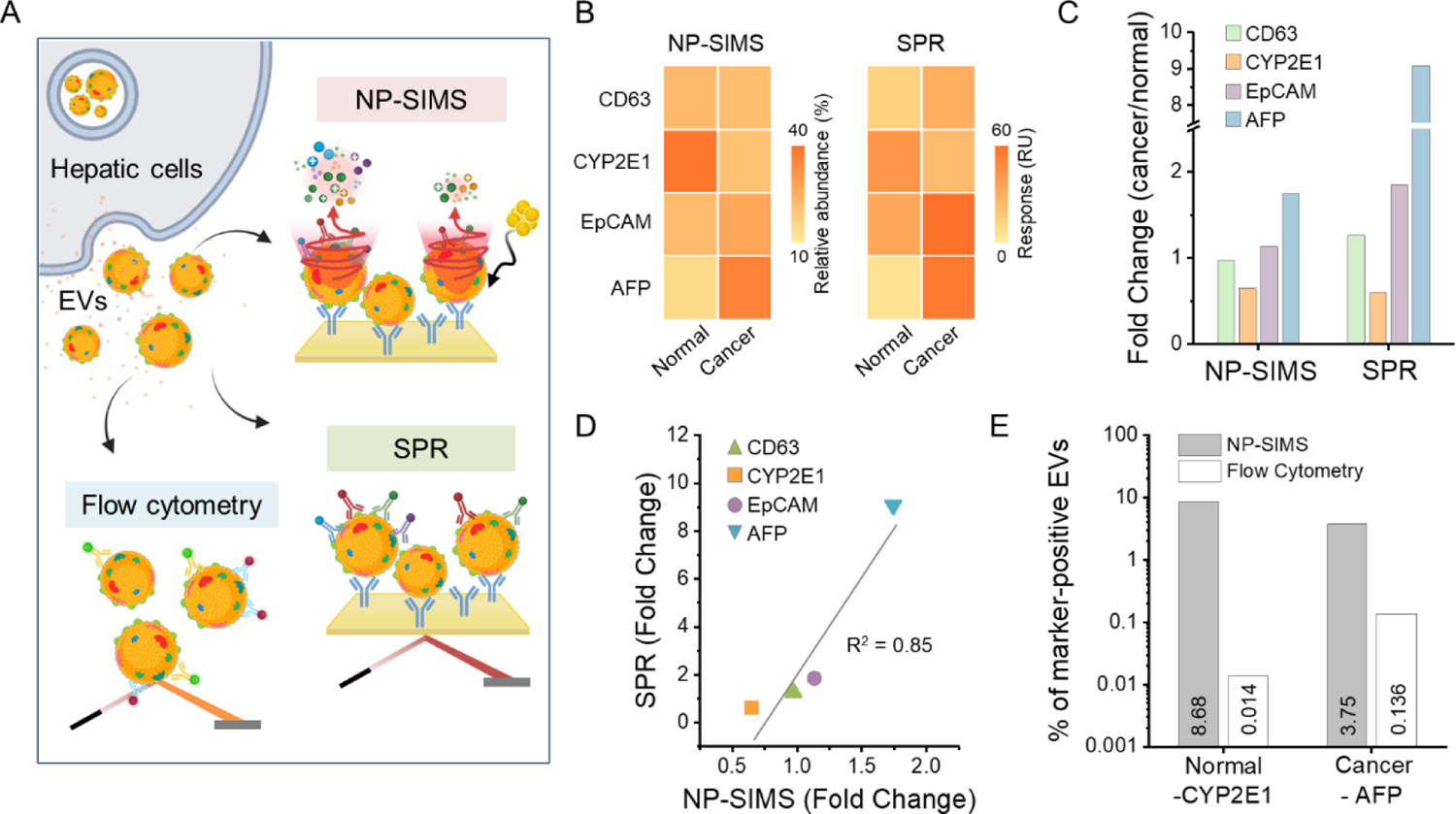
Benchmarking NP-SIMS against SPR and nanoflow cytometry. (A) EVs were harvested from media conditioned by normal (primary human hepatocytes) and cancer (HepG2) cells. The same EV sample was split three ways and analyzed by NP-SIMS, SPR and nanoflow cytometry. All four markers were analyzed by NP-SIMS and SPR, two markers (CYP2E1 and AFP) were analyzed by nanoflow cytometry. (B) Heatmap comparing of surface marker expression on normal and cancer EVs for NP-SIMS and SPR. (C) Comparison of EV surface marker expression determined by NP-SIMS and SPR. Fold change values for NP-SIMS were calculated based on relative abundance measurements. (D) Regression analysis showing close agreement between two approaches. (E) Head-to-head comparison of NP-SIMS and nanoflow cytometry. %CYP2E1-posoitive normal EVs and %AFP-positive cancer EVs were used for differentiating the two methods. All of the results were obtained with three biological replicates (n=3).

SPR represents an excellent benchmarking technology for several reasons: (1) it uses substrates (Au-coated chips) and Ab immobilization chemistry that is identical to NP-SIMS, (2) SPR is a sensitive, quantitative, and label-free means of monitoring molecular binding events. We carried out SPR analysis of the same set of EV samples analyzed by NP-SIMS.

Figure 4B presents EV marker expression in a heatmap format. The left side of the heatmap shows the relative abundance (%) values measured by NP-SIMS while the right side presents response units (RUs) from SPR measurements. One may appreciate that the two methods yielded similar results for EV surface marker expression. Another way to compare the two analytical approaches was to quantify the fold change of biomarker expression on cancer and normal EVs for SPR and NP-SIMS. As may be seen from Figure 4C**,D**, the two analytical approaches were in general agreement, with the exception of higher AFP levels measured by SPR.

Next, we carried out our comparison of NP-SIMS with nanoflow cytometry using the same set of EVs. Because our nanoflow cytometer was limited to two biomarkers (fluorophores), we chose to focus on CYP2E1 and AFP for analysis of EVs from normal and cancer liver cells. The nanoflow cytometry results, presented in **Figure S11**, may be used to draw several conclusions. First, nanoflow cytometry supported our previous observations with NP-SIMS that AFP is more prevalent in EVs derived from HepG2 cells. Second, AFP-positive EVs were ∼9 fold more frequent compared to CYP2E1-positive EVs in the sample derived from HepG2 cells, which is also in alignment with NP-SIMS. However, smaller (2-fold) differences in AFP and CYP2E1 expression in normal EVs seen in NP-SIMS and SPR data were not captured by nanoflow cytometry. This may be explained by the fact that nanoflow cytometry reports on frequency of detection events and not on the levels of expression. Therefore, it may not be sensitive enough for scenarios where changes in expression levels are small and do not translate into a different frequency of detection events. NP-SIMS is unique in that it quantifies the relative abundance of biomarkers on a given EV and may be used to group individual EVs based on relative levels of biomarker expression.

Another observation is the differences in sensitivity of NP-SIMS and nanoflow cytometry (see Figure 4E). Out of 10^10^ events (particles/mL) counted by the nanoflow cytometer, only 10^6^ (0.01%) were identified as positive based on the fluorescence signal for CYP2E1 or AFP. In contrast, using NP-SIMS to examine the sample containing a 1:1 mixture, 5000 out of 79000 (5%) EVs were categorized as cancer or normal EVs based on the detection of two Ab-MS tags, anti-CD63@^150^Nd and either anti-CYP2E1@^142^Nd or anti-AFP@^161^Dy. This indicates that NP-SIMS ∼500 times more sensitive than nanoflow cytometry for analyzing EVs. Overall, benchmarking studies support the accuracy of NP-SIMS analysis of EV composition and highlight benefits of our approach – single EV resolution, high sensitivity, and quantitation of the biomarker expression levels.

## CONCLUSIONS

In this paper, we describe the use of NP-SIMS in conjunction with Abs carrying Ln-tags for detection of surface markers on individual EVs. The NP-SIMS approach allowed analysis of thousands of individual EVs for the presence and abundance of surface markers. Simultaneous detection of up to four EV surface markers was demonstrated. Importantly, results of NP-SIMS analysis were rigorously benchmarked using established technologies, SPR and nanoflow cytometry. To the best of our knowledge, this is the first report of mass spectrometry analysis of individual EVs using Ln-tags. Our approach holds considerable promise for multiplexed analysis of large numbers (thousands to millions) of individual EVs to identify EV subsets based on the presence, levels, and co-localization of surface markers. As a step toward this goal, we demonstrated that the levels of hepatic surfaces markers, CYP2E1 and AFP, may be used to differentiate EVs derived from normal and malignant hepatocytes. Furthermore, NP-SIMS was used to accurately determine composition in a sample containing a mixture of EVs and revealed information about co-localization of molecules on a nanometer scale. It is important to note that ∼40 Ln tags are currently available commercially and that the NP-SIMS technology is compatible with these tags. This means that multiplexing capabilities of our approach may be expanded in the future. In summary, NP-SIMS represents a platform technology for multiplexed analysis of thousands of single EVs and may be used in the future for profiling EV-based biomarkers of liver cancer and other diseases.

## MATERIALS AND METHODS

### Materials

4-morpholineethanesulfonic acid (MES), MUA, EDC, NHS, and sodium azide were purchased from Sigma-Aldrich (St. Louis, MO, USA). Dulbecco’s modified Eagle’s medium (DMEM), fetal bovine serum (FBS), penicillin/streptomycin were purchased from Gibco (Grand Island, NY, USA). Ethyl alcohol (EtOH) was purchased from Electron Microscopy Sciences (Hat eld, PA, USA), while isopropyl alcohol (IPA) was purchased from Honeywell (Charlotte, NC, USA). Exosome-Depleted FBS was purchased from Captivate Bio (Watertown, MA, USA). Total cell culture EV isolation kit, EV spin column (MW 3000), Alexa Fluor™ 488 Antibody Labeling Kit, and Alexa Fluor™ 647 Antibody Labeling Kit were purchased from Invitrogen (Carlsbad, CA, USA). Dulbecco’s Phosphate-buffered Saline (DPBS) was purchased from Corning (Corning, NY, USA). Mouse anti-human CD63 and mouse IgG isotype control were purchased from BD Biosciences (San Jose, CA, USA). Rabbit anti-human CYP2E1 Ab was purchased from CYP450-GP (Vista, CA, USA). Rabbit IgG isotype control and Vybrant DiO cell-labeling solution were purchased from Thermo Fisher Scientific (Waltham, MA, USA). Anti-EpCAM and Anti-AFP were purchased from R&D Systems (Minneapolis, MN, USA). Maxpar® X8 Antibody Labeling Kit was purchased from Fluidigm (South San Francisco, CA, USA). Antibody stabilizer solution was purchased from Boca Scientific (Dedham, MA, USA).

### Cell Culture

Human hepatocytes and HepG2 cells were used for EV harvesting and NP-SIMS analysis. Human hepatocytes were a kind gift from Prof. Takeshi Saito at University of South California and Phoenix Bio Ltd. These cells were isolated from chimeric mice with humanized livers (cDNA-uPA+/−/SCID (uPA+/wt: B6;129SvEv-Plau, SCID: C.B-17/Icr-scid/scid Jcl)) using standard cannulation and collagenase perfusion protocol^50–52^ and cultured in DMEM supplemented with 20 mM HEPES, 10% FBS, 100 U/mL penicillin, 100 μg/mL streptomycin, 15 μg/mL L-Proline, 0.25 μg/mL recombinant human insulin, 50 nM Dexamethasone, 5 μg/mL EGF, 0.1 mM L-ascorbic acid 2-phosphate, 2% DMSO at 37 °C in a humidified 5% CO_2_ atmosphere. HepG2 cells were passaged and cultured in DMEM supplemented with 10% FBS, 100 U/mL penicillin, 100 μg/mL streptomycin at 37 °C in a humidified 5% CO_2_ atmosphere. For EV harvesting, the cells were washed twice with PBS to eliminate EVs derived from FBS and then cultured in EV-depleted DMEM supplemented with 5% EV-depleted FBS for 72 h.

### Isolation and Characterization of EVs

EVs were isolated using a total EV isolation kit. First, the cell culture media from ∼10^7^ cells was collected and subjected to centrifugation at 2,000g for 30 min at 4 °C to remove cellular debris. Then, the EV isolation reagent was added to the supernatant and allowed to incubate for 16 h at 4 °C. Subsequently, the EVs were extracted by subjecting the mixture to centrifugation at 10,000g for 1 h at 4 °C, and then resuspended in 1× PBS and stored at −80 °C until use. For fluorescence labeling, EVs were incubated with Vybrant DiO cell labeling solution for 30 min at 37 °C. The excess dye was then removed from the EVs using Exosome Spin Columns. The size distribution and concentration of isolated EVs were measured via NTA (Malvern Panalytical, Malvern, U.K.). The EVs were captured on substrates identical to those used for NP-SIMS analysis (Au surface functionalized with anti-CD63) and were examined using a Hitachi S-4700 cold field emission SEM (Hitachi High Technologies America, Inc., Schaumburg, IL).

### Preparation of the Abs@Ln-Tag

Maxpar Labeling Kits were used to create metal-isotope-Ab (Abs@Ln tag) complexes. For multiplexed detection, four different Abs specific to target proteins were conjugated to MS tags selected for their high probability of ionization and detection - anti-CD63@^150^Nd, anti-CYP2E1@^142^Nd, anti-EpCAM@^146^Nd, and anti-AFP@^161^Dy. In addition, isotype or negative control Abs (mouse IgG) were also conjugated with ^150^Nd tag following the manufacturer’s instructions (i.e. anti-IgG@^150^Nd). The Abs@Ln tags were redispersed in antibody stabilizer solution with 0.05% sodium azide to obtain a final concentration of 0.5 mg/mL of conjugated antibody and stored at 4 °C for further use.

### NP-SIMS Analysis of EVs

Pieces of silicon wafers (10 by 10 mm^2^) with a 100 nm layer of sputtered Au were used as substrates for NP-SIMS analysis.^41^ The Au surfaces were functionalized with anti-CD63 Abs using a workflow described by us previously. ^42, 53^ Briefly, Au-coated Si wafers were cleaned by sonication in IPA for 30 min, then exposed to oxygen plasma for 5 min at 150 mW (YES-G500, Yield Engineering Systems, Freemont, CA, USA). Cleaned Au substrates were first treated with MUA (10 mM in pure EtOH) for 12 h to create a self-assembled monolayer of alkanethiol with terminal carboxylic acid groups. After SAM formation, the substrates were thoroughly rinsed with ethanol and deionized (DI) water, and dried using nitrogen. Next, the carboxylic groups of MUA were activated with EDC/NHS mixture (a 1:1 ratio of 200 mM EDC and 100 mM NHS in 0.1 M MES buffer) for 1 h to create amine-reactive moieties. The surfaces were rinsed with DI water, and the substrates were immediately incubated with an anti-CD63 solution (50 µg/mL in PBS) for 1.5 h. The substrates were then washed with 1× PBS to remove excess Abs and blocked by incubating for 1 h in BSA (1% in 1× PBS) to prevent nonspecific binding. After washing with 1× PBS again, functionalized substrates were incubated with the EV sample (10^9^ particles/mL in 1× PBS) for 2 h and then thoroughly washed with 1× PBS. When detecting multiple surface markers, substrates with captured EVs were exposed sequentially to four target-specific MS tags - anti-CD63@^150^Nd, anti-CYP2E1@^142^Nd, anti-EpCAM@^146^Nd, and anti-AFP@^161^Dy. All Ab-tag conjugates were at a concentration of 5 µg/mL and were incubated for 1 h. A substrate was washed thoroughly with 1xPBS between incubation with individual Ab-tag conjugates. The EV samples were prepared at Mayo Clinic, were rinsed with DI water, dried using nitrogen and shipped on dry ice to Texas A&M University for NP-SIMS analysis. Upon receipt, the samples were brought up to room temperature in a desiccator and mounted into the NP-SIMS sample holder using carbon tape. The samples were then analyzed using a custom-built NP-SIMS instrument described by us previously.^54^ Briefly, this instrument utilizes a liquid metal ion source to produce a range of Au cluster projectiles. A Wien filter is used to select Au_400_^4+^ projectiles and to accelerate these projectiles toward the target achieving a final kinetic energy of 414 keV. After each projectile impact, the resulting SIs were extracted and analyzed with a reflectron time-of-flight mass spectrometer. The instrument is operated in the event-by-event bombardment/detection mode, meaning that SIs emitted from each impact are resolved and analyzed. Data analysis was carried out using in-house mass spectrometry software.^30^

### SPR Analysis of EVs

SPR experiments were conducted using a three-channel system (Biosensing Instruments, Tempe, AZ). Au SPR chips were prepared by self-assembly of MUA, activation of carboxylic groups with EDC− NHS, and immobilization of anti-CD63 Abs. BSA was then used for blocking the surface to minimize nonspecific bonding. The functionalization steps and concentration of reagents/sample were the same as described for NP-SIMS analysis above. The MUA-treated Au chips were mounted into the SPR instrument and then used immediately for anti-CD63 functionalization. The flow rate used to inject reagents in the SPR instrument was 20 μL/min. EVs were flowed in the SPR at 10^9^ particles/mL in 1× PBS and at the flow rate of 10 µL/min for 10 min. Then the surface of the chip was flushed with running buffer (1× PBS) for 8 min, and the baseline for SPR measurement was established. Subsequently, Abs@Ln (5 µg/mL in PBS) were perfused over captured EVs at a flow rate of 20 µL/min for 5 min. Afterwards, the SPR chip was flushed with running buffer for 8 min and the final baseline was recorded. SPR response (resonance units or RU) was determined by monitoring the changes in baseline before and after the sample injection.

### Nonoflow Cytometry Analysis of EVs

EVs were analyzed using an Apogee A60 Micro Plus System (A60MP, Apogee Flow Systems Inc., Northwood, UK) as previously described.^28, 29^ Briefly, anti-CYP2E1 and anti-AFP Abs were conjugated with Alexa Fluor 488 (AF488) and Alexa Fluor 647 (AF647) antibody labeling kits (Thermo Fisher Scientific, Waltham, MA), respectively. For sample preparation, 5 μL of EV samples (10^10^ particles/mL) were incubated with 10 μL of anti-CYP2E1@AF488 and anti-AFP@AF647 with desired concentrations (18 μg/mL in PBS for both antibodies). Following incubation in the dark for 30 min at room temperature, the antibody-EVs mixture proceeded to sample analysis. A blank sample with 1× PBS was examined before each sample analysis to ensure a count rate < 100 events per second. Each sample was analyzed three replicates for 60 seconds. Number of detected events, sample dilution (200 for this sample), flow rate (1.5 μL/min) and acquisition time (60 s) were used to determine EV particle concentration (particles/mL). Data analysis was performed using FlowJo version 10.6.1.

## ASSOCIATED CONTENT

### Supporting Information

SPR analysis to characterize EV capture and labeling; SPR analysis of hepatic EVs; NP-SIMS analysis to characterize multiplexing detection; Characterization of hepatic EVs using SEM, fluorescence microscopy, and NTA; Characterizing EV heterogeneity; Methodology for computing Colocalization Factor values; Nanoflow cytometry analysis of EVs.

## AUTHOR INFORMATION

### Corresponding Author

E-mail (Emile A. Schweikert; Alexander Revzin; Michael J. Eller).

### Author Contributions

S.L., D.S.V., S.V.V., K.G., and Y.K., performed the experiments; S.L., D.S.V., M.J.E., S.V.V., M.A.S., Y.K., F.L., H.M., A.R., and E.A.S. analyzed the data; S.L., D.S.V., M.J.E., S.V.V., A.R., and E.A.S. wrote the manuscript with input from all authors. All authors have given approval to the final version of the manuscript.

### Notes

The authors declare no competing financial interest.

## ACKNOWLEDGMENTS

This work was funded by grants from NIH (GM123757 and CA236612). Additional support was provided by Cells to Cures Strategic Initiative at Mayo Clinic.

## Table of Contents

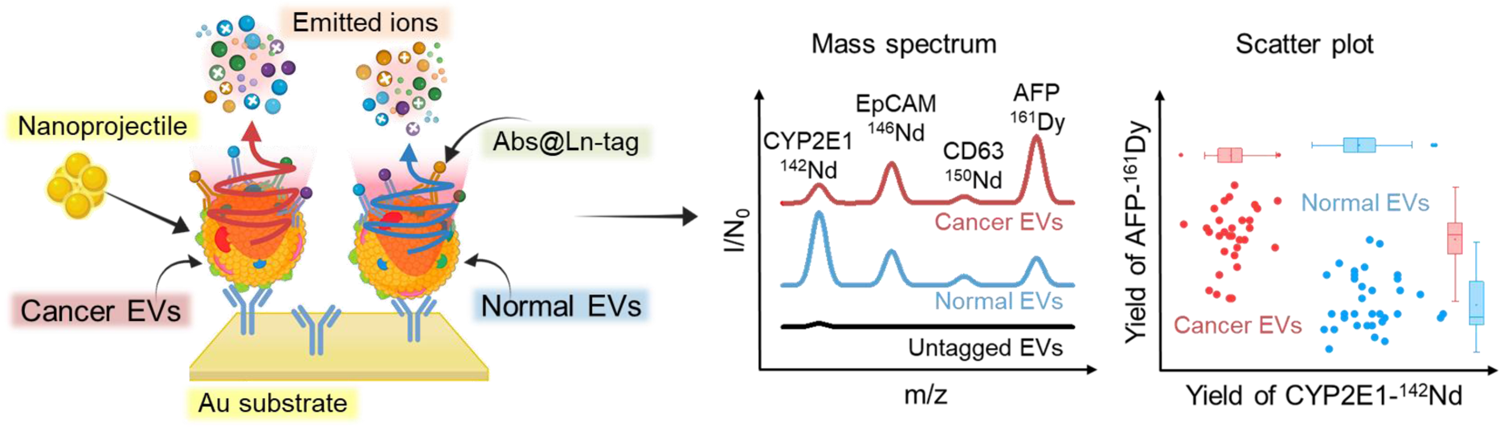

