## Supporting information for "Nanoprojectile Secondary Ion Mass Spectrometry Enables Multiplexed Analysis of Individual Hepatic Extracellular Vesicles"

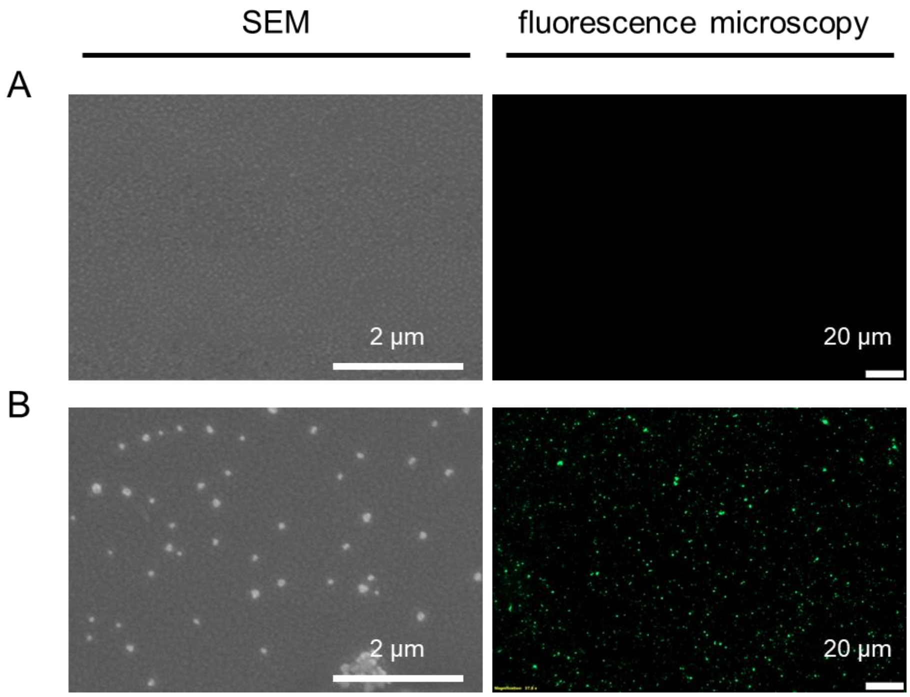


**Figure S1**. **SEM and fluorescence microscopy images of EVs on substrates.** EVs were labeled with DiO dye and captured on Au substrates functionalized with anti-CD63. (A) Images of the surface before and (B) after incubating with solution of EVs ($10^9$ particles/mL).

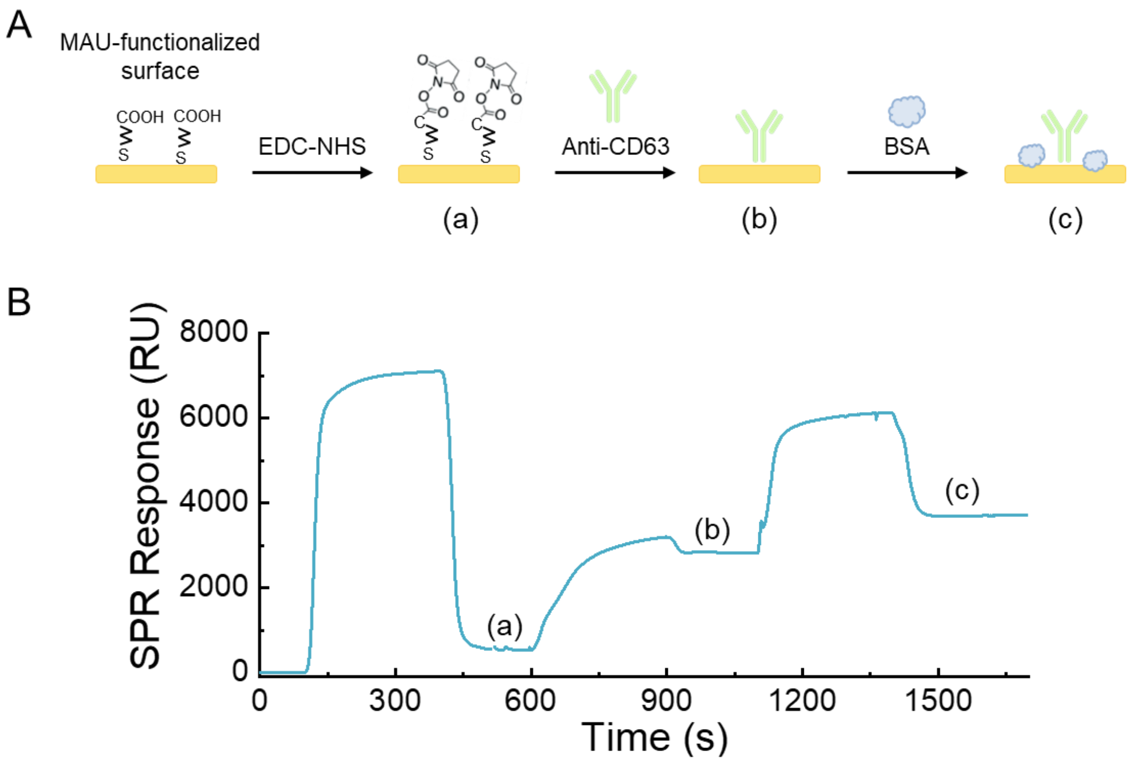


**Figure S2. Description of SPR characterization of surface modification steps.** (A) Au substrates were functionalized by assembly of MUA layer, making this layer amine-reactive by treatment with EDC/NHS, then conjugating anti-CD63 Abs, and blocking with BSA. Reagents for each step were introduced sequentially into SPR chip. (B) SPR signals associated with each step in the surface functionalization process. RU: Resonance Units.

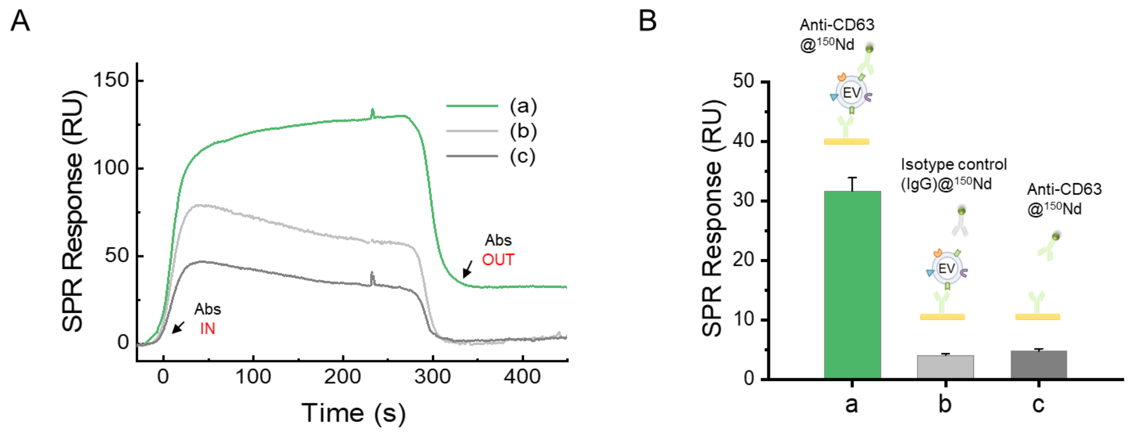


**Figure S3**. **Assessing specificity of interactions between EVs and Ab@Ln tags using SPR.** (A) SPR response curves three experiments scenarios: (a) EVs captured on the surface and then challenged with anti-CD63@$^{150}$Nd tags, (b) EVs captured on the surface and then challenged with isotype control (anti-IgG@$^{150}$Nd) tags and (c) Au substrate surface without EVs challenged with anti-CD63@$^{150}$Nd tags. (B) Comparison of SPR results highlights specificity of interaction between EVs and anti-CD63@$^{150}$Nd tags. Signals were similar to noise for interactions between EVs and isotype control tags or for tags interacting with surfaces without EVs. Data are displayed as mean ± SD (n=3). RU: Resonance Units.

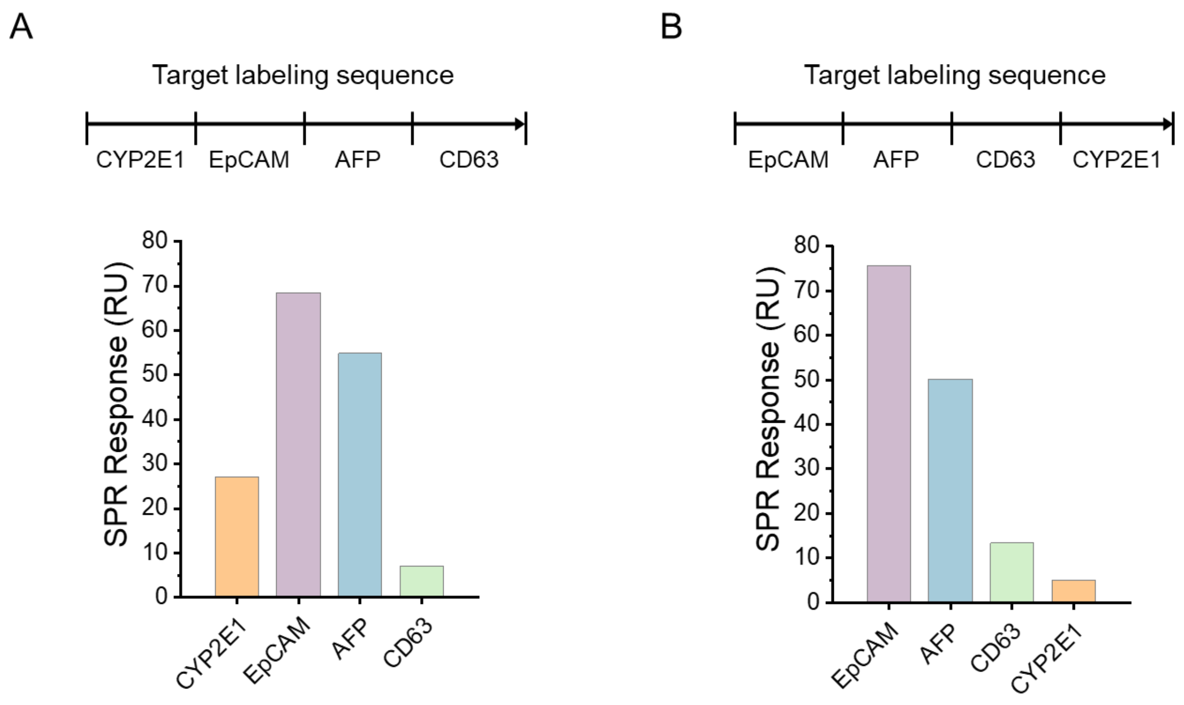


**Figure S4**. **Evaluating sequence of labeling steps using SPR analysis.** (A,B) Ab@Ln tags were introduced individually into SPR flow cells. Two labeling sequences were tested. RU: Resonance Units.

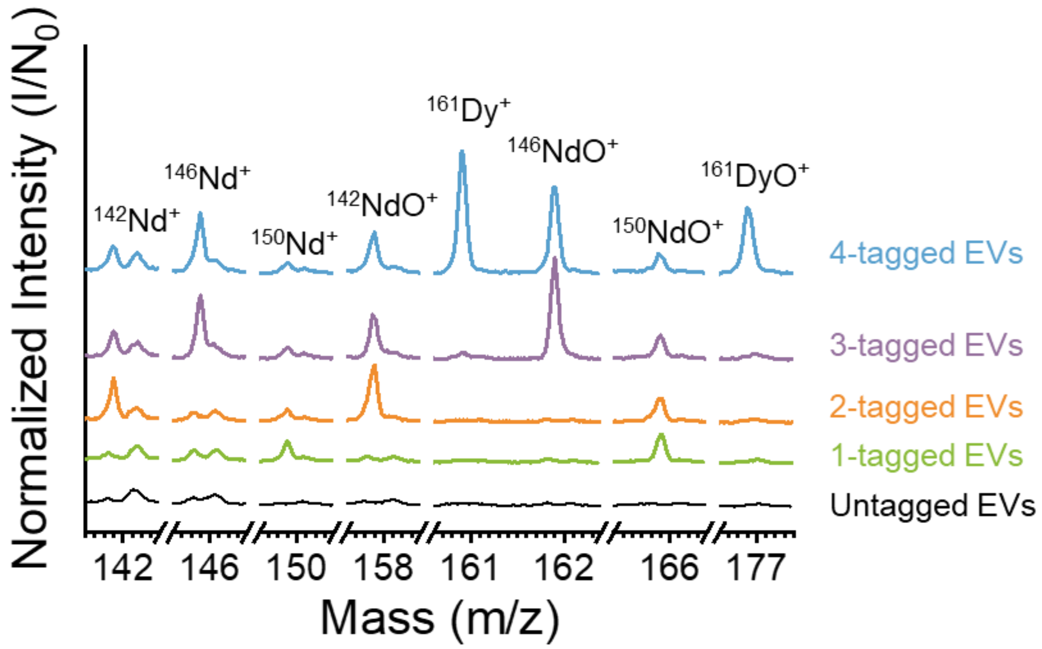


**Figure S5. Characterizing multiplexed analysis of EVs using NP-SIMS.** Selected regions of mass spectra comparing several EV tagging scenarios. Multiple characteristic ions associated with four MS tags originating from EVs ($^{150}Nd^{+}$ and $^{150}NdO^{+}$ for CD63, $^{142}Nd^{+}$ and $^{142}NdO^{+}$ for CYP2E1, $^{146}Nd^{+}$ and $^{146}NdO^{+}$ for EpCAM and $^{161}Dy^{+}$ and $^{161}DyO^{+}$ for AFP).

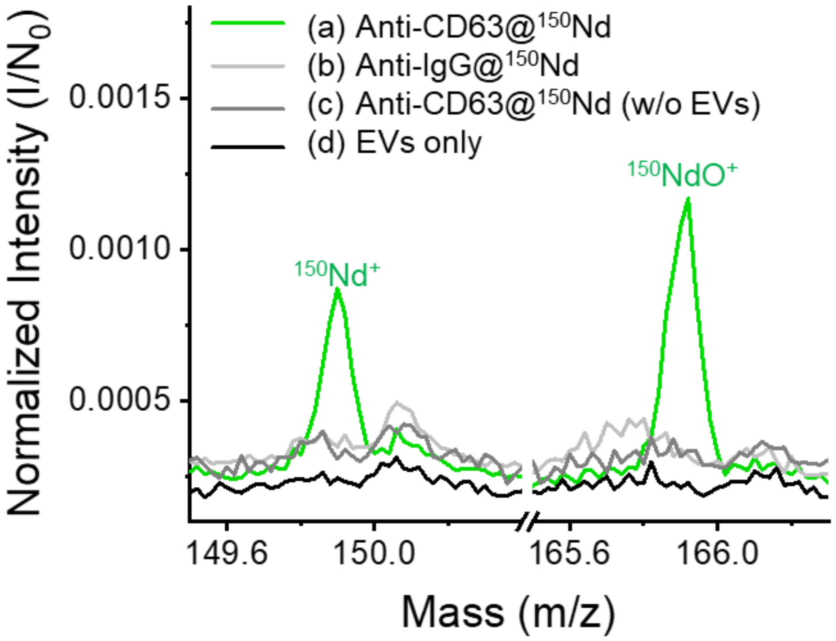


**Figure S6**. **Assessing specificity of Abs@Ln tag – EV interactions with NP-SIMS.** Comparison of peak intensity ($I/N_0$) of (a) Anti-CD63@$^{150}$Nd and (b) Anti-IgG@$^{150}$Nd to captured EVs and (c) anti-CD63@$^{150}$Nd to EV free surface. The data are binned to the nearest 0.02 m/z for comparison.

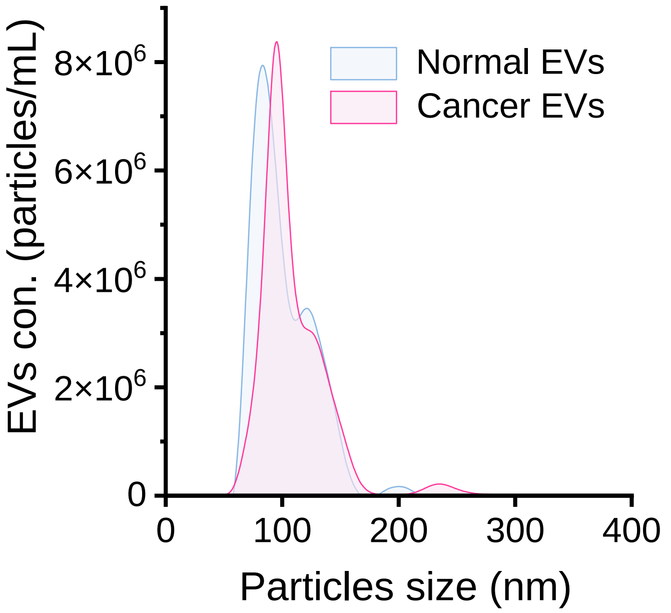


**Figure S7. NTA analysis of EVs harvested normal (human hepatocytes) and cancer (HepG2) cells.** The normal EVs exhibited a mean diameter of 112 ± 21 nm and a concentration of $2.35 \times 10^{10}$ particles/mL, while the cancer EVs showed a mean diameter of 119 ± 28 nm and a concentration of $3.48 \times 10^{10}$ particles/mL.

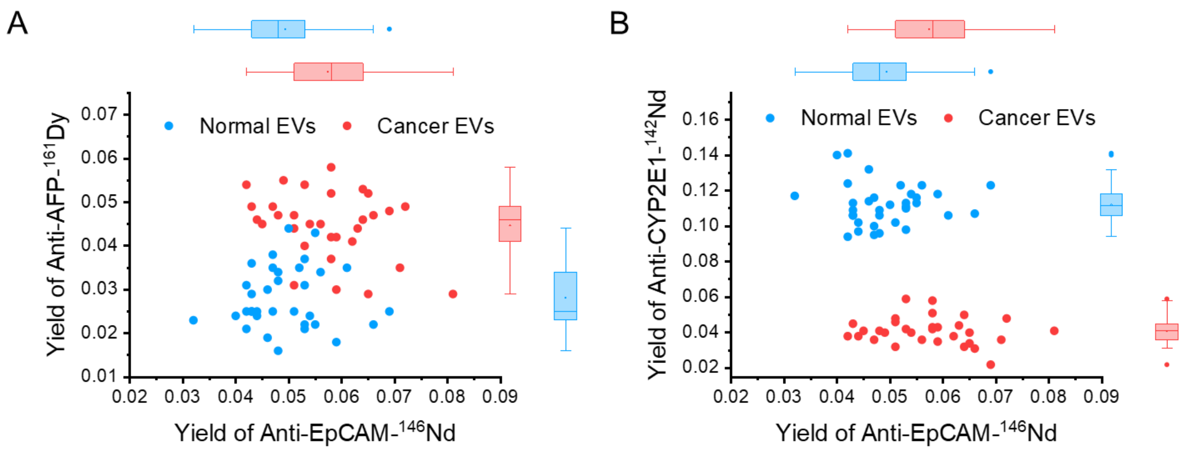


**Figure S8**. **Analysis of heterogeneity in EV biomarker expression.** EVs were harvested from either normal hepatocytes or cancer cells. EVs from each sample were captured on individual substrates, analyzed by NP-SIMS. CD63 gating was used to assign signals as emanating from EVs. (A) AFP versus EpCAM and (B) CYP2E1 versus EpCAM scatter plots. Each point corresponds to 1000 EVs of each type.

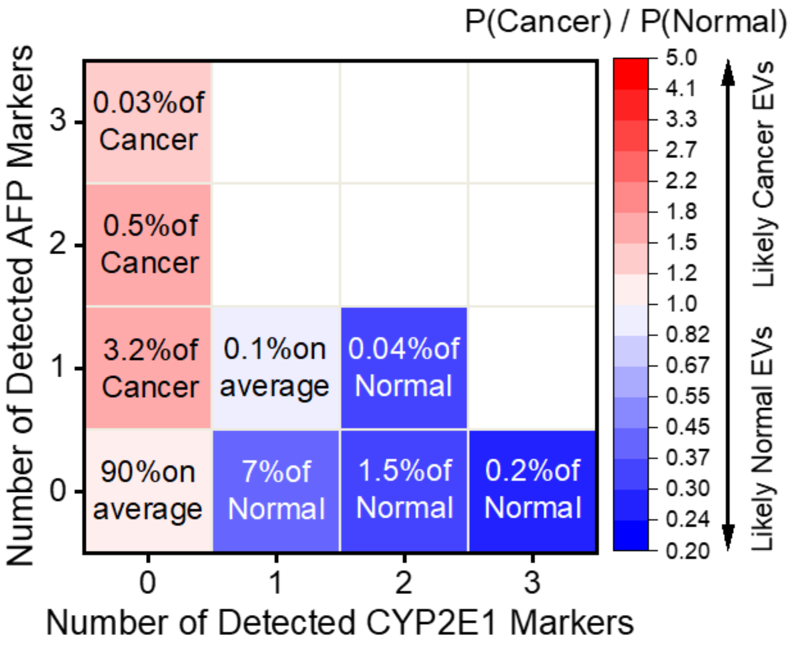


**Figure S9. Comparison of mass spectrometry signals from cancer and normal EVs.** Data on individual EVs are binned based on the number of detected AFP tag ions (0, 1, 2, 3 AFP) and the number of detected CYP2E1 tag ions (0, 1, 2, 3 CYP2E1). The probabilities of each of these conditions are compared (cancer/normal) where values greater than unity are characteristic of cancer EVs and values less than unity are characteristic of EVs from normal cells.

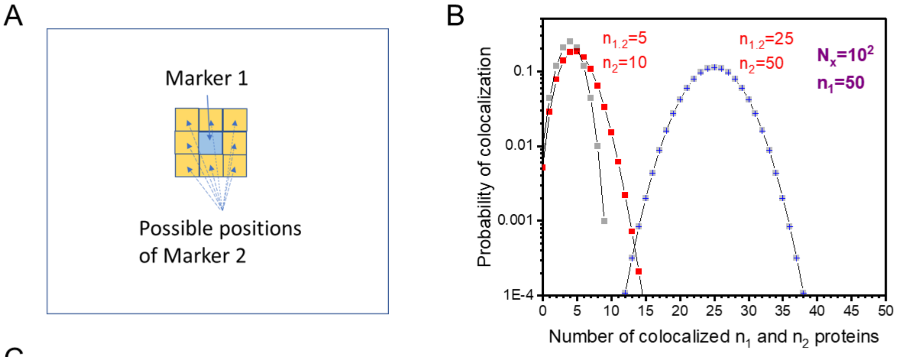


C

Measured number of markers

| Type of EVs | $N_{CD63}$ | $N_{CYP2E1}$ | $N_{EpCAM}$ | $N_{AFP}$ | $\sum N_i$ |
|---|---|---|---|---|---|
| Normal EVs | $1.6 \times 10^5$ | $6.5 \times 10^5$ | $3.8 \times 10^5$ | $2.6 \times 10^5$ | $1.45 \times 10^6$ |
| Cancer EVs | $1.5 \times 10^5$ | $2.7 \times 10^5$ | $5.5 \times 10^5$ | $6.7 \times 10^5$ | $1.64 \times 10^6$ |

D

Relative colocalization factors, *C*

| Cancer EVs | | | | |
|---|---|---|---|---|
| | CYP2E1 | EpCAM | CD63 | AFP |
| CYP2E1 | | 0.69 | 0.20 | 1.34 |
| EpCAM | | | 0.29 | 2.08 |
| CD63 | | | | 10.99 |
| AFP | | | | |

| Mix cancer / normal EVs 1:1 ratio | | | | |
|---|---|---|---|---|
| | CYP2E1 | EpCAM | CD63 | AFP |
| CYP2E1 | | 0.50 | 1.92 | 0.77 |
| EpCAM | | | 1.14 | 0.76 |
| CD63 | | | | 4.82 |
| AFP | | | | |

**Figure S10. Evaluating colocalization of molecules on EVs.** (A) Sketch of co-positions of markers. (B) Dependence of probability of random colocalization on the number of colocalized tagged proteins (markers), $n_{1,2}$. (C) Number of markers of each type measured for EVs from human hepatocytes (normal) and HepG2 (cancerous) cells. To compare the number of colocalized markers, the Colocalization Factor is used. (D) Relative colocalization factors, *C*, for any pair combinations of four markers. The accuracy of values of the relative colocalization factors is <5%.

*1. Model of random colocalization (theoretical approach – occupancy theory)*

Let's consider a sub-ensemble of EVs where each EV has an exact number $n_1$ and $n_2$ of the tagged proteins (markers) of two types. One should note that for this sub-ensemble, the number of randomly colocalized markers, $n_{1,2}$ , is the probability function, $P(n_1,n_2)$.[1]

$$P(n_{1,2}) = \alpha \frac{n_2!}{n_{1,2}!(n_2-n_{1,2})!} \left(\frac{n_1}{n_x}\right)^{n_{1,2}} \left(1-\frac{n_1}{n_x}\right)^{n_2-n_{1,2}} \qquad (1)$$

where the $n_x$ is the number of the available areas of size of the marker, at the surface of an EV, $\alpha$ is the geometry factor ($\alpha \leq 8$). The geometry factor reflects the variety of co-positions of markers at the surface plane (**Figure S10A**). **Figure S10B** shows the dependence of $P(n_1,n_2)$ on number of colocalized markers. This dependence shows that the number of randomly colocalized markers is small for the expected number of markers (tens of each type) at the surface of an EV (EV diameter ~120 nm).

To compare the model with an experimental data, the average number of colocalized markers, $\overline{n_{1,2}}$, is computed for the probability distribution (1).

$$\overline{n_{1,2}} = \propto \frac{n_1 n_2}{n_x} \qquad (2)$$

This dependence of $\overline{n_{1,2}}$ for randomly colocalized markers shows a linear dependence on $n_1$ and $n_2$. If the colocalization is not random, this dependence should show a nonlinear behavior with respect to the number of markers. This is the case discussed below for the cancer EVs.

*2. Single cluster projectile impact on colocalized tagged proteins (markers)*

The samples made for the NP-SIMS experiments are discussed in the main text (Paragraph: Comparison of EVs from normal and cancer cells). These samples (normal, cancer, and 1:1 normal-cancer EVs) have been bombarded with ~1 keV/atom $Au_{400}$ projectiles to obtain the collections of individual mass spectra (~ $10^6$ spectra) for each sample. The analysis of the individual mass spectra allows to compute: a) the number of detected tag ions (e.g. $Nd^+$ and $NdO^+$), $I_n$ ($n \leq 4$), b) the number of co-detected tag ions, $I_{n,m}$ ($n,m \leq 4$), which are linked to the colocalized markers.

The single impact on the surface of an $Au_{400}$ projectile stimulates emission from ~ 10 nm size area. This emission process follows two significant statistical features: a) statistical equivalency of impacts on physically/chemically indistinct surface areas, and b) independence of emissions of secondary ions from such areas.[2] Thus, for the impacts on the areas where the markers are colocalized, the yield of emitted ions (number of emitted/detected per impact ions), $Y_{1,2}$, is

$$Y_{1,2} = \frac{x}{\beta} Y_1 Y_2 \qquad (3)$$

where $Y_1$ and $Y_2$ are the yields of the emitted tag ions from marker 1 and marker 2. The geometry factor, $\beta \leq 4$, shows that for projectile impacts on both markers, the yield for the most probable impact factor is less ~2 times for each marker. One should note that the geometry factor is the same for any pair of markers. The yields are: $Y_1 = \frac{I_1}{N_1}$ (4) and $Y_2 = \frac{I_2}{N_2}$ (5) , and $Y_{1,2} = \frac{I_{1,2}}{N_{1,2}}$ (6), where $I_1$, $I_2$ and $I_{1,2}$ are the number of detected ions, which were emitted from the markers 1, 2, and the colocalized markers 1 and 2 respectively. Using the expressions (3)-(6), one can compute the relationship of the number of colocalized markers, $N_{1,2}$, and the total number of markers, $N_1$, $N_2$.

$$N_{1,2} = \frac{\beta I_{1,2}}{I_1 I_2} N_1 N_2 \qquad (7)$$

The $N_{1,2}$ from (7) depends on the measured values solely. The expression (7) is similar to the model expression (3), except that instead of the parameter of the random colocalization, $n_x$, the expression (7) contains the effective correlation coefficients, $N_x = \frac{I_1 I_2}{I_{1,2}}$. This coefficient may indicate the enhancement of colocalization toward the random model case. The measured number of markers of each type is presented in the **Figure S10C**.

*3. Computing colocalization factors*

The Colocalization Factor, $K_{1,2}$, is the normalized number of colocalized markers. For instance, for the pair of markers CD63 (general marker) and AFP (cancer marker), the $K_{\mathrm{CD63,AFP}}$ is

$$K_{\mathrm{CD63,AFP}} = \frac{N_{\mathrm{CD63,AFP}}}{\sum_{i=1,..4} N_i}$$

where the $\sum N_i$ is the sum of all detected markers of four types.

The colocalization factors can be measured for any pair of different types of markers. To compare the colocalization factors measured for normal and cancer EV samples, the relative colocalization factors are used.

$$C = \frac{K^{(c)}_{\mathrm{CYP2E1,AFP}}}{K^{(n)}_{\mathrm{CYP2E1,AFP}}}$$

where the superscripts (n) and (c) are the symbols which indicate normal and cancer EV samples. The **Figure S10D** shows the relative colocalization factors for the sample of cancer EVs, and the sample of the mixture (1:1) of cancer and normal EVs. The factor $C$=11 **(Figure S10D)**, measured for the pair of the cancer marker AFP and the general marker CD63, shows that the number of AFP markers is not only increased at the surface of cancer EVs (**Figure

**S10C**), but also, these markers experienced the effect of enhancement above the random colocalization (expression (2)) with the general CD63 markers. In addition, for the sample of 1:1 of cancer and normal EVs, the factor CD63,AFP is half of this factor measured for the sample of 100% of cancer EVs.

(1) Riordan, J. Introduction to combinatorial analysis; John Wiley & Sons Inc.: New York, 1958; pp 90.

(2) Rickman, R.; Verkhoturov, S.; Parilis, E.; Schweikert, E. Simultaneous ejection of two molecular ions from keV gold atomic and polyatomic projectile impacts. Phys. Rev. Lett. 2004, 92 (4), 047601.

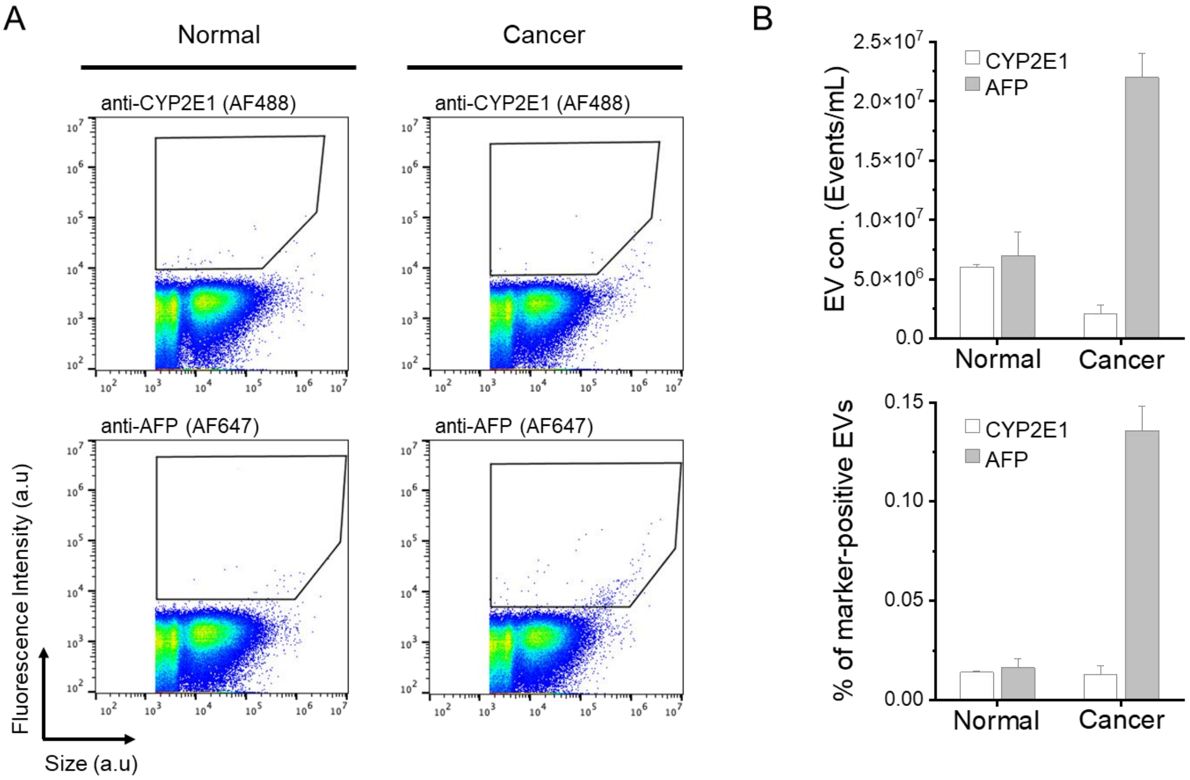


**Figure S11**. **Nanoflow cytometry analysis of EV samples.** (A) Representative nanoflow cytometry analysis of CYP2E1 and AFP expression in normal and cancer EVs. Scatter plots from of cancer EVs show higher level of AFP expression. (B) Quantification of CYP2E1 and AFP events in normal and cancer EVs. EV concentration (top) and marker-positive particles (bottom). The marker-positive particles (%) are calculated by normalized to total particle concentrations.
